# Contribution of PSD-95 protein to reward location memory

**DOI:** 10.1101/590109

**Authors:** Anna Cały, Małgorzata Alicja Śliwińska, Magdalena Ziółkowska, Kacper Łukasiewicz, Roberto Pagano, Agata Nowacka, Malgorzata Borczyk, Kasia Radwanska

## Abstract

The molecular mechanisms involved in formation of memory are still poorly understood. We focus here on the function of post-synaptic density protein 95 (PSD-95) and its phosphorylation by CaMKII in spontaneous learning about reward location in female mice. We show that formation of reward location memory leads to downregulation of PSD-95 protein in dendritic spines of the *stratum radiatum*, area CA1, and selective shrinkage of dendritic spines that contain PSD-95. ShRNA-driven, long-term downregulation of PSD-95 in the area CA1 decreases precision of memory. Autophosphorylation deficient CaMKII mutant mice (CaMKII:T286A) need more time than wild-type animals to learn the location of reward. The same impairment is observed after CA1-targeted overexpression of CaMKII phosphorylation-deficient form of PSD-95 (PSD-95:S73A). In contrast to young adult mice, in aged animals reward location learning affects only spines that lack PSD-95. The frequency and size of the spines without PSD-95 are increased, while shRNA targeted to PSD-95 affects neither speed of learning nor precision of memory indicating alternative mechanisms to support successful memory formation in old mice. Altogether, our data suggest that dynamic regulation of PSD-95 expression is a mechanism that accelerates learning and improves precision of reward location memory in young mice. The function of PSD-95 in memory processes changes in aged animals.

## Introduction

The ability to find food and other natural rewards, as well as remember their location, is a key to animal survival. In humans, the importance of this process can be appreciated when perception of reward is aberrant, leading to exaggerated and inflexible reward seeking in drug addiction. Therefore, understanding molecular and cellular basis of reward seeking and memory of its location is crucial to understand processes involved in affective disorder such as addiction or depression.

Formation and consolidation of memory involves functional and structural plasticity of excitatory synapses ^1–3^. Post-synaptic density protein 95 (PSD-95/SAP90), a member of the membrane-associated guanylate kinase (MAGUK) family, is highly abundant in the post-synaptic density (PSD) of an excitatory synapse and has been proposed to regulate different forms of synaptic transmission ^4–10^, synapse structure and stability ^11–13^ as well as formation and long-term stabilisation of memory ^14–17^. PSD-95-dependent protein complexes interact both with AMPA-and NMDA-type glutamate receptors (AMPARs and NMDARs), and PSD-95 regulates NMDAR-dependent changes in AMPARs number ^18–21^. Synaptic localization and function of PSD-95 is controlled by many interacting proteins and modifications, including phosphorylation, palmitoylation and ubiquitination ^9,22–27^. In particular, upon stimulation of NMDAR, calcium and calmodulin-dependent kinase II (CaMKII)-driven phosphorylation of PSD-95 at serine 73 (PSD-95: S73) controls interactions of PSD-95 with NMDAR, synaptic localization of PSD-95, growth of dendritic spine and synaptic plasticity ^26,28^. Still, the role of dynamic regulation of PSD-95 protein in the synapse is poorly understood in the context of memory process.

This study sought to understand the contribution of PSD-95 to reward location memory by integrating *ex vivo* analysis of PSD-95 expression, virally-mediated manipulation of PSD-95 expression and mobility, as well as behavioural analysis. We demonstrate that overexpression of CaMKII phosphorylation-deficient PSD-95 (PSD-95:S73A) in the area CA1 slows-down learning about reward location, while depletion of PSD-95 levels by shRNA impairs precision of memories. This process operates in young mice, and is impaired in aged animals what may underlie age-related cognitive decline.

### Animals

αCaMKII autophosphorylation-deficient mutant mice (αCaMKII-T286A) ^29^, and heterozygous of Thy1-GFP M line mice (Thy1-GFP ^+/-^) ^30^ were bred (as heterozygotes with the 129J/C57BL/6J background) in the Animal House of the Nencki Institute of Experimental Biology, and genotyped as previously described ^29,30^. Young, adult mice were 5±1 month-old during the behavioral training, whereas old individuals were 20±2 month-old. Only female mice were used for all experiments, as male are too aggressive for group housing in the IntelliCages. All mice were housed with access to food and water *ad libitum*, and 12:12 hour dark-light cycle, 23–24°C and 35-45% humidity. The studies were carried out in accordance with the European Communities Council Directive of 24 November 1986 (86/609/EEC), Animal Protection Act of Poland and approved by the 1st Local Ethics Committee in Warsaw. All efforts were made to minimize the number of animals used and their suffering.

### Reward location memory test in IntelliCages

The IntelliCage system (NewBehavior AG, Zürich, Switzerland) (http://www.newbehavior.com/) consists of a large standard rat cage (20.5 cm high, 40 cm × 58 cm at the top, 55 cm × 37.5 cm at the base). In each corner, a triangular learning chamber is located with two bottles. To drink, only one mouse can go inside a plastic ring (outer ring: 50 mm diameter; inner ring: 30 mm diameter; 20 mm depth into outer ring) that ends with two 13 mm holes (one on the left, one on the right) that provides access to bottle nipples. Each visit to the corner, nosepoke at the doors governing access to the bottles, and lick were recorded by the system and ascribed to a particular animal. During experiments in each cage two corners were active. Groups of 8 to 15 mice were housed per cage.

Mice were subcutaneously injected with unique microtransponders which allow for mice identification in the IntelliCage (10.9 mm length, 1.6 mm diameter; Datamars, Slim Microchip T-SL) under brief isoflurane anesthesia. Animals were allowed to recover for 3 days after the injection and after this time they were introduced to the IntelliCage. Experiments consisted of two phases: habituation (8-12 days) and learning. During habituation, animals had access to water in all corners. Four or five days before the learning phase, mice got access to 5% sucrose solution (in tap water) from the top of the cage to get familiarized with its taste. Baseline corner preference was measured during the last day of the habituation as the % of visits or licks. During learning, water in less preferred corner was replaced by 5% sucrose. The change in preference for the corner with sucrose (% visits performed to sucrose corner versus all visits) during training, compared with the preference of the same corner during the baseline period (H – the last day of habituation) was used as an index of spatial learning. All phases of the training were started at the beginning of the dark phase (12:00 a.m.).

### Immunostaining on brain slices

Mice were anesthetized and transcardially perfused with filtered PBS (Sigma–Aldrich) followed by 4% PFA (Sigma–Aldrich) in PBS. Brains were removed and placed overnight in the same fixing solution and afterwards in 30% sucrose in PBS for three days. Next, coronal brain sections (40 μm thick) were prepared (Cryostat Leica CM1950, Leica Biosystems Nussloch GmbH, Wetzlar, Germany) and stored at –20 °C in PBSAF [PBS, 15% sucrose (Sigma-Aldrich), 30% ethylene glycol (Sigma-Aldrich), and 0.05% NaN_3_ (SigmaAldrich)]. The sections were washed with PBS, PBS/0.3%/Triton X-100 (Sigma-Aldrich) followed by 1-h incubation in a blocking solution (5% normal donkey serum in PBS/0.3% Triton X-100) and overnight incubation with the antibodies directed against PSD-95 (1:500, MAB1598; Merck-Millipore, RRID:AB_94278). Next, the sections were washed in PBS with 0.3% Triton X-100 and incubated for 90 minutes with the secondary antibody: anti-mouse Alexa Fluor 555 (1:500, A31570, Invitrogen, RRID:AB_2536180). The sections were mounted on glass microscope slides (Thermo Fisher Scientific), air-dried and coverslipped with Fluoromount-G medium with DAPI for fluorescence (00-4959-52, Invitrogen).

The staining was analyzed with the aid of confocal, laser-scanning microscope. Z-stacks of dendrites in the CA1 were acquired using Zeiss Spinning Disc microscope (63× oil objective and 1.66 digital magnification) (Zeiss, Göttingen, Germany). A series of 18 continuous optical sections (67,72 µm × 67,72 µm), at 0.26 μm intervals along the *z-*axis of the tissue section, were scanned. Six to eight Z-stacks of microphotographs were taken per animal, from every sixth section through the dorsal hippocampus (*stratum radiatum* of CA1 field) (one dendrite per neuron per image). Z-stacks were reconstructed to maximal projections and analyzed with ImageJ software. Threshold tool was used, which identifies objects distinct from the background based on intensity. The density and average size of PSD-95+ puncta, as well as their co-localization were analyzed using Fiji software and measured using the analyze particle tool as previously described ^31^. To analyze the images of the stained sections with overexpression of AAVs, a confocal microscope (magnification: x63, oil objective) (Leica TCS SP8, Leica Microsystems, Wetzlar, Germany) was used, and mean gray value of the microphotographs was assessed with ImageJ software.

Dendritic spines filled with GFP (in Thy1-GFP mice) were analyzed using semiautomatic SpineMagick! Software ^32^. Data analysis was performed using scripts in Python. Overall we analyzed: 1112 spines from young, control mice; 2455 spines from young, learning mice; 972 spines from old, control animals and 1367 spines from old training group. Custom-written Python scripts were used for Fiji software to analyze co-localization of PSD-95+ puncta with dendritic spines.

### Stereotactic intracranial injections

Mice were anaesthetized with isoflurane (5% for induction, 1.5-2.0% after), fixed in the stereotactic frame (51503, Stoelting, Wood Dale, IL, USA), and their body temperatures were maintained using a heating pad. Stereotactic injections were performed bilaterally into CA1 region of hippocampus using coordinates from the Bregma: AP, −2.1mm; ML, ±1.1 mm; DV, −1.3mm according to ^33^. 0.5 µl of virus solution was microinjected through beveled 26 gauge metal needle and 10 µl microsyringe (SGE010RNS, WPI, USA) connected to a microsyringe pump (UMP3, WPI, Sarasota, USA), and its controller (Micro4, WPI, Sarasota, USA) at a rate 0.1 µl/min. The microsyringe was left in a place for additional 10 min following injection to prevent leakage of the vector. Mice were injected with AAV1/2 coding wild-type form of PSD-95 (AAV:αCaMKII-PSD95(WT)-mCherry-WPRE) (0.5 µl/ site, viral titer 1,35 ×10^9^/μl), the mutated form of PSD-95 with point substitution of serine 73 to alanine (AAV:αCaMKII-PSD95(S73A)-mCherry-WPRE) (0.5 µl/ site, viral titer 9,12 ×10^9^/μl), or control mCherry (AAV:αCaMKII-mCherry-WPRE (0.5 µl/ site, viral titer 7,5 ×10^7^/μl, obtained from Deisseroth’s Lab). Lentiviral vectors (LVs) coding short-hairpin RNA silencing PSD-95 expression (αCaMKII-shRNA(PSD95)-GFP (0.5 µl/ site, viral titer 2,52 ×10^8^/μl) (gift from Dr. Oliver M. Schlüter (European Neuroscience Institute Göttingen, Germany) ^8^ or control vector based on a pSUPER shRNA targeting the *Renilla* luciferase cloned into pTRIP (H1-shRNA(luciferase)) (0.5 µl/ site, viral titer 6,52 ×10^8^/μl) (donated by Dr Katarzyna Kalita, Nencki Institute of Experimental Biology, Warsaw, Poland) were used. The viruses were prepared by Animal Model Core Facility at Nencki Institute.

After the surgery, animals were allowed to recover for 14 days before the training in the IntelliCages. After the training the animals were perfused with 4%PFA in PBS and Zeiss Spinning Disc confocal microscope (magnification: 10×) was used to photograph the dorsal hippocampus and assess the extent of the viral expression.

### Statistical data analysis

Data acquisition and quantification was performed in a group blind manner. All statistical analyses were performed using Prism 6 (GraphPad Software). The exact sample size (e.g., the number of mice or spines) of each experiment is provided in the relevant figures together with details of statistical tests. For behavioural data, immunostaining and dendritic spine analysis one-way and two-way analysis of variance (ANOVA), and post-hoc Tukey’s multiple comparisons test were used. Dendritic spine volume did not follow normal distributions and were compared with Mann-Whitney test. For other parameters, unless specified, t-tests were performed. All data with normal distribution are presented as the means ± standard error of the mean (SEM). For samples which did not follow normal distribution medians and interquartile range (IQR) are shown. The difference between the experimental groups was considered as significant if p < 0.05.

## Results

### Formation of memory about reward location downregulates PSD-95 protein in dendritic spines

To study neuronal processes underlying learning of reward location we used IntelliCage setup. In this setup the activity and spontaneous learning of female mice leaving in a group can be measured in close to ecologic conditions and without stressful intrusion of the experimentators ^34^. We used young adult Thy1-GFP(M) mice (5±1 month-old) ^30^ (**Fig. 1A**) to analyse co-localisation of PSD-95 protein and dendritic spines as a proxy of training-induced synaptic remodelling ^35,36^. Mice were trained to find sucrose reward in one of two active cage corners ^34^ (**Fig. 1B.i**). Animals significantly increased preference of the rewarded corner during the first 30 minutes of the training, and continue to prefer this corner during the following 90 minutes (**Fig. 1B.ii**).

**Figure 1.**
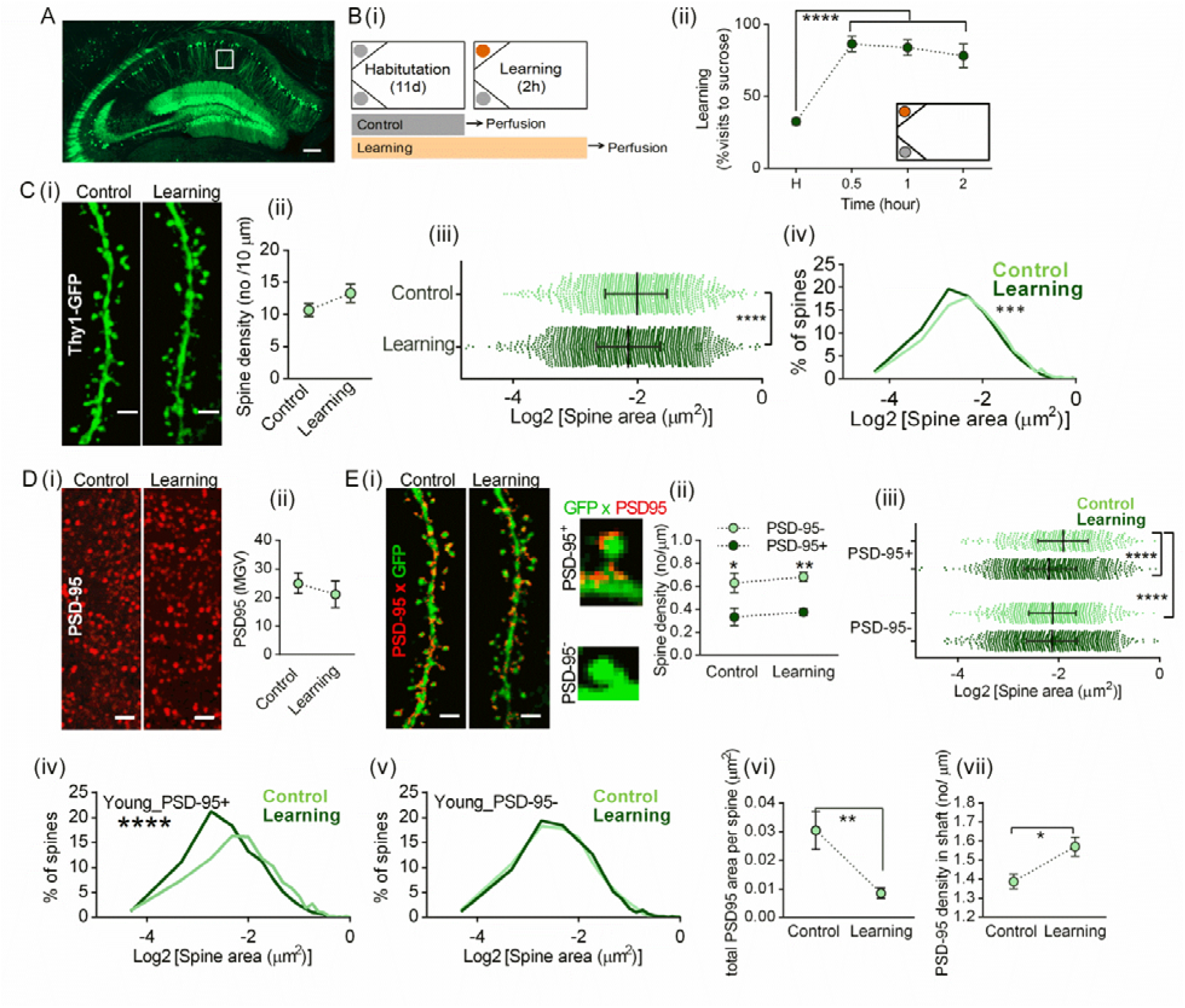
Learning-induced remodeling of PSD-95 protein scaffolding and dendritic spines. **(A)** Representative image of dorsal hippocampus of Thy1-GFP(M) mouse. A white rectangle indicates the area (*stratum radiatum* of dorsal area CA1) were dendritic spines and protein expression were analyzed. Scale bar: 200 μm. **(B) (i)** Cage setups during training and experimental timeline. The cages had two active corners during training: with water (gray circle) and sucrose (orange circle). **(ii)** Mean+/- SEM preference of the reward corner during training (control mice, n = 5; learning, n = 6). Mice increased preference of the reward corner during the first 30 minutes of the training (****p<0.0001, RM ANOVA with Tukey’s multiple comparisons test). H, preference of the corner during the last day of the habituation. Inset, cage setup during training. **(C) (i)** Representative high resolution images of dendritic fragments of the control and trained Thy1-GFP(M) mice, scale bars: 2 µm. **(ii)** Linear density of dendritic spines was not affected by the training (t-test, t(9) = 1.447, p > 0.05). **(iii)** Dendritic spines shrank during training (*****p < 0.0001, Mann-Whitney test, U = 1236337). Graph shows data for individual spines (control, mice/ spines = 4/ 1112; learning, mice/ spines = 5/ 2455) with medians and interquartile range. Scales are Log_2_-tranformed. All statistical tests are performed on non-transformed data. (**iv**) Distributions of the spine sizes in learning mice shifted from control to smaller sizes (***p=0.0004, Kolmogorov-Smirnov test, D = 0.07417). **(D)** (**i**) Representative high resolution images of PSD-95 immunostaining in control and trained mice, scale bars: 2 µm. (**ii**) Mean gray value of the images was not affected by the training (p > 0.05, t-test, t(9) = 0.800; control, n = 5 and trained mice, n = 6). **(E) (i)** Representative high resolution images of PSD-95 immunostaining co-localized with dendritic fragments of the control and trained Thy1-GFP(M) mice, scale bars: 2 µm. Right, spines with PSD-95 (PSD-95+) and without PSD-95 (PSD-95-). (**ii**) Average+/-SEM density of the spines. Spines without PSD-95 are more frequent than spines with PSD-95. Training does not affect density of the spines (*p<0.05, **p<0.001 PSD-95-vs PSD-95+, two-way ANOVA with Tukey’s multiple comparisons test) (control PSD-95-, mice/ spines = 4/ 613; learning PSD-95-, mice/ spines = 5/ 1430; control PSD-95+, mice/ spines = 4/ 440; learning PSD-95+, mice/ spines = 5/ 1020). (**iii**) In control mice PSD-95+ spines are bigger than PSD-95-spines (****p<0.0001). Only PSD-95+ spines shrank after training (****p<0.0001, Kruskal-Wallis ANOVA followed by Dunn’s multiple comparisons tests, H = 35.18). (**iv**) Distribution of PSD-95+ spines’ areas shifted from control to smaller values after training (****p<0.0001, Kolmogorov-Smirnov test, D = 0.1501). **(v)** The distribution of PSD-95-spines’ areas did not change after learning (D = 0.02615). **(vi)** Average area of PSD-95 immunostaining puncta per spine decreased after training (control, n = 4; learning = 6) (**p < 0.01, t-test, t(8) = 3.826). (**vii**) Average density of PSD-95 puncta in dendritic shaft increased after training (control, n = 4; learning = 6) (*p < 0.05, t-test, t(8) = 2.927).

Next, we analysed dendritic spines in *stratum radiatum* of CA1 area (**Fig. 1C.i**), as this region is involve in formation of spatial memory ^37^. Training did not affect density of spines (**Fig. 1C.ii**). However, median dendritic spines’ areas were smaller after training as compared to control mice (**Fig. 1C.iii)**, and distribution of dendritic spines’ areas was shifted to smaller values after learning as compared to the spines analysed in control mice (**Fig. 1C.iv)**.

To study the expression of PSD-95 protein we performed immunostaining with PSD-95-specific antibody and analysed its co-localisation with dendritic spines. Intensity of PSD-95 immunostaining in the area CA1 was not changed in the learning mice as compared to the control group (**Fig. 1D**). When dendritic spines were segregated in two categories: with and without PSD-95 [PSD-95(+) and (-)] (**Fig. 1E.i**), we observed that only 43% of the spines contained PSD-95, which is very low as compared with previous studies showing that in the visual cortex (V1) over 80% of spines contained PSD-95 protein ^38^. Therefore, to validate our method, we analysed PSD-95 protein expression in dendritic spines of the V1 cortex in young control animals. The frequency of dendritic spines with PSD-95 in V1 reached 80%, as previously reported ^38^. The frequency of PSD-95-positive spines in the same animals in the area CA1 was 45%. The spines in V1 were also bigger, and contained more PSD-95 puncta as compared to CA1 region (extended data **Fig. 1-1**). Thus, although we cannot exclude the possibility that we did not detect PSD-95 protein if it was expressed in very low quantity, we concluded that low frequency of dendritic spines that contain PSD-95 protein plausibly indicates low frequency of mature spines in the area CA1 ^39,40^, as compared to the cortical region.

We subsequently calculated density and size of dendritic spines with and without PSD-95 after training to find that the mean densities of the spines of these two categories were not affected by the training, and PSD-95(-) spines were more frequent than PSD-95(+) spines after learning, as in the control animals (**Fig. 1E.ii**). The analysis of the areas of spines showed that the spines with PSD-95 have higher median values than spines without PSD-95 (**Fig. 1E.iii**). Moreover, the median of PSD-95(+) spines’ areas decreased after training while the median of PSD-95(-) spines’ areas was not changed (**Fig. 1E.iii**). The change of PSD-95(+) spines was also observed as a shift of size distribution toward smaller values in learning mice as compared to the controls (**Fig. 1E.vi**). No change in distribution of spines’ areas was observed in PSD-95(-) spines (**Fig. 1E.v**). We also analysed PSD-95 puncta to find that the total area of PSD-95 puncta per PSD-95(+) spines was decreased in the learning group, as compared with the controls (**Fig. 1E.vi**), while density of PSD-95 puncta in the shaft increased (**Fig. 1E.vii**) suggesting translocation of the protein.

In summary, our data indicate remodelling of dendritic spines during memory formation that is dendritic spine type-specific. In young adult mice, training to locate sucrose reward results in shrinkage of big spines containing PSD-95 in the area CA1. At the same time PSD-95 protein level in dendritic spines is decreased and the protein is partly translocated to the shaft.

### PSD-95 regulates precision of reward location memory

To test the function of PSD-95 protein in reward location memory, we used lentiviruses encoding short hairpin RNA (shRNA) targeted to PSD-95 mRNA (LV:αCaMKII-shRNA_PSD-95-GFP) ^8^ (**Fig. 2A**). Four-month old, C57BL/6J mice had LVs stereotactically injected into dorsal area CA1 and 14 days later they were trained in the IntelliCages (**Fig. 2A.i-ii**). ShRNA for PSD-95 effectively knocked down the endogenous PSD-95 protein in the area CA1 (39% decrease), as compared with the control virus coding shRNA designed for *Renilla* luciferase (LV:H1-shRNA_luciferase) (**Fig. 2A.iii-iv**). ShRNA for PSD-95 did not impair mice performance during initial 30 minutes of the training, however, later the preference of the reward corner of the mice transfected with shRNA for PSD-95 was lower than the preference of the control animals (**Fig. 2AB.v**).

**Figure 2.**
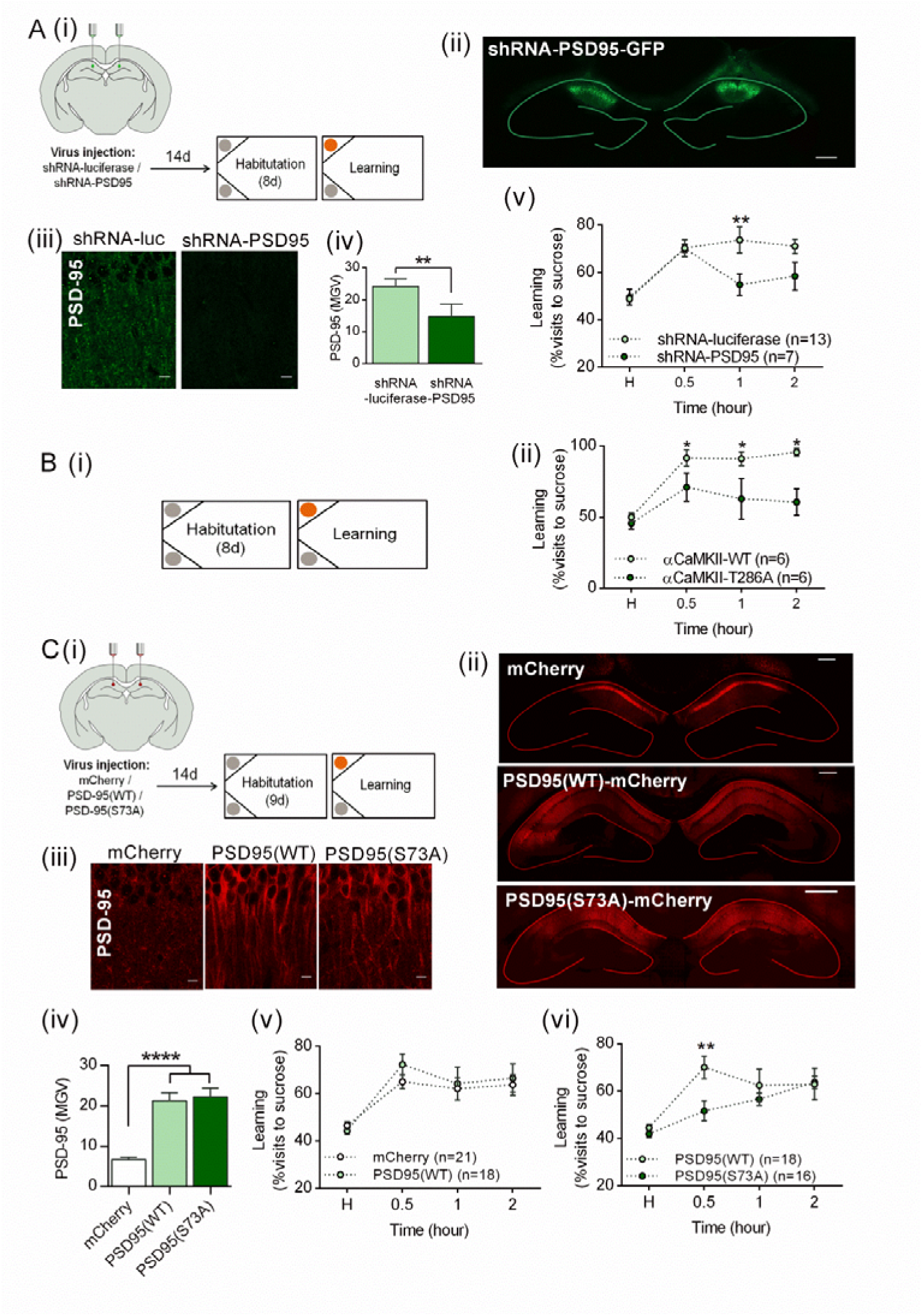
PSD-95 protein in the area CA1 controls spatial learning. **(A) (i)** Experimental timeline. (**ii**) Representative microphotography of dorsal hippocampus with local expression of lentiviruses coding shRNA targeted to PSD-95 (LV:αCaMKII_shRNA PSD-95_GFP). Scale bar: 100 µm. **(iii)** Microphotographs of PSD-95 immunostaining in the *stratum radiatum* of dorsal area CA1 after local expression of shRNA to *Renilla* luciferase (LV:αCaMKII_shRNA_luc_GFP) or PSD-95 (LV:αCaMKII_shRNA_PSD-95_GFP). Scale bars: 10 µm. (**iv)** shRNA for PSD-95 decreased mean gray value of PSD-95 immunostaining (**p = 0.0054, t-test, t(6) = 4.243; mice with shRNA-luc: n = 5; shRNA-PSD95: n = 3). **(v)** ShRNA for PSD-95 impaired preference for reward corner after 1 and 2 hours of the training (**p < 0.01, Two-way RM ANOVA with Sidak’s multiple comparisons test, virus: F_1, 76_ = 12.72, p = 0.0006, time: F_7, 76_ = 8.611, p < 0.0001). H, preference of the corner during the last day of the habituation. **(B) (i)** Experimental timeline of spatial training of αCaMKII-T286A mutant mice and their WT littermates. **(ii)** T286A mutants are impaired in formation of spatial memory (*p < 0.5, two-way ANOVA with Sidak’s multiple comparisons test, genotype: F_1, 72_ = 11.02, p = 0.001, time: F_7, 72_ = 5.173, p < 0.0001). **(C) (i)** Experimental timeline. (**ii**) Representative images showing bilateral expression of AAV1/2 coding PSD95(WT)_mCherry, PSD95(S73A)_mCherry or mCherry in the area CA1. Scale bars: 100 µm. **(iii)** Representative microphotographs of the area CA1 showing PSD-95 protein immunostaining after viral infection. Scale bars: 10 µm. (**iv**) Transfection of CA1 area with AAVs coding PSD95(WT)_mCherry, PSD95(S73A)_mCherry resulted in overexpression of PSD-95 protein (**** p < 0.0001, one-way ANOVA by Tukey’s multiple comparisons test, virus: F_2, 22_ = 24.30, p < 0.0001). **(v)** Overexpression of PSD-95(WT) did not affect preference for the reward corner (Two-way RM ANOVA with Sidak’s multiple comparisons test, virus: F_1, 36_ = 0.5392, p = 0.4675; time: F_3, 108_ = 12.53, p < 0.0001), (**vi**) while overexpression of phosphorylation deficient PSD-95(S73A) decreased preference for the rewarded corner during the first 0.5 hr of the training (**p<0.01, Two-way RM ANOVA with Sidak’s multiple comparisons test, virus: F_1, 30_ = 3.139, p = 0.0866; time: F_3, 90_ = 8.053, p < 0.0001).

In summary, our experiments indicate that long-term downregulation of PSD-95 protein in CA1 does not affect formation of reward location memory but results in poor precision of memory, as demonstrated by long-term decrease of the reward corner preference.

### Autophosphorylation of CaMKII and CaMKII-dependent phosphorylation of PSD-95 regulates speed of learning

Synaptic localization of PSD-95 is controlled by interacting proteins and post-translational modifications. In particular, upon stimulation of NMDAR, calcium and calmodulin-dependent kinase II (CaMKII)-driven phosphorylation of PSD-95 at serine 73 (PSD-95: S73) controls interactions of PSD-95 with NMDAR, synaptic localization of PSD-95, growth of dendritic spine and synaptic plasticity ^26,28^. We therefore decided to test whether CaMKII-dependent phosphorylation of PSD-95:S73 controls reward location memory.

First, to test the role of CaMKII in spatial memory formation, we used 4-month old, autophosphorylation-deficient αCaMKII mutant mice (αCaMKII-T286A) ^29^. Autophosphorylation of CaMKII: threonine 286 decreases clustering of PSD-95 with NMDAR subunit, NR2B ^26^. The young T286A mutants, as compared with the young WT mice, had decreased preference of the reward corner during training indicating impaired formation and precision of spatial memory (**Fig. 2A**).

To test the role of the interaction of CaMKII with PSD-95 protein in spatial memory formation we used AAV1/2, coding wild-type (WT) and phosphorylation-deficient mutant PSD-95 protein at CaMKII-targeted Serine 73 (S73A) ^26,28^. In control group we used AAV1/2 coding mCherry. The viruses were stereotactically injected into dorsal area CA1 of 4-month old, C57BL/6J mice (**Fig. 2C.i**), resulting in overexpression of PSD-95 protein (**Fig. 2C.ii and iii**). Two weeks after the surgery mice were trained (**Fig. 2C.i**). Overexpression of PSD-95(WT) did not affect the preference of the reward corner during training, as compared to mCherry control (**Fig. 2C.v**). Overexpression of phosphorylation-deficient mutant PSD-95(S73A), as compared to wild-type form of PSD-95, decreased preference of the reward corner during initial 30 minutes of training, but not at the later time points (**Fig. 2C.vi**). Since overexpression of PSD-95 in CA1 and CaMKII-T286A mutation affected general activity of the mice (extended data, **Fig. 2-1**), we also analysed the preference of the reward corner in 10-visit bins (to make the number of learning trials equal between the experimental groups) to find similar effects as described for the timebins (extended data, **Fig. 2-1**).

In summary, our experiments indicate that phosphorylation of PSD-95 at serine 73 accelerates memory formation during initial 30 minutes of the training, while autophosphorylation of CaMKII is controls both speed and long-term precision of memory in young mice.

### Regulation of PSD-95 expression during reward location training is age dependent

A growing body of evidence indicates that during ageing many synaptic processes in the hippocampus are impaired (Burke and Barnes 2006, 2010), presumably leading to compromised precision of spatial memory and speed of learning (Hedden et al. 2004). We therefore asked whether learning-induced remodelling of PSD-95 protein at the synapse is altered in aged mice.

We compared morphology of dendritic spines and expression of PSD-95 protein in young adult (5±1 month-old) and old (20±2 month-old) Thy1-GFP(M) mice trained in the IntelliCages (**Fig. 3A**). Young mice were more active than old animals during the habituation, but not during the training when both groups of mice increased frequency of visits (extended data, **Fig. 3-1.A**). Both young and old mice significantly increased preference of the rewarded corner during the first 30 minutes of the training (**Fig. 3A**), up to circa 85%, and there was no statistically significant difference between old and young mice (**Fig. 3A).** Similar pattern of the preference for reward corner was observed when licks were analysed (extended data, **Fig. 3-1.A**), or visit preference in 10-visit bins (extended data, **Fig. 3-1.A**), suggesting no gross cognitive impairment in old Thy1-GFP(M) mice.

**Figure 3.**
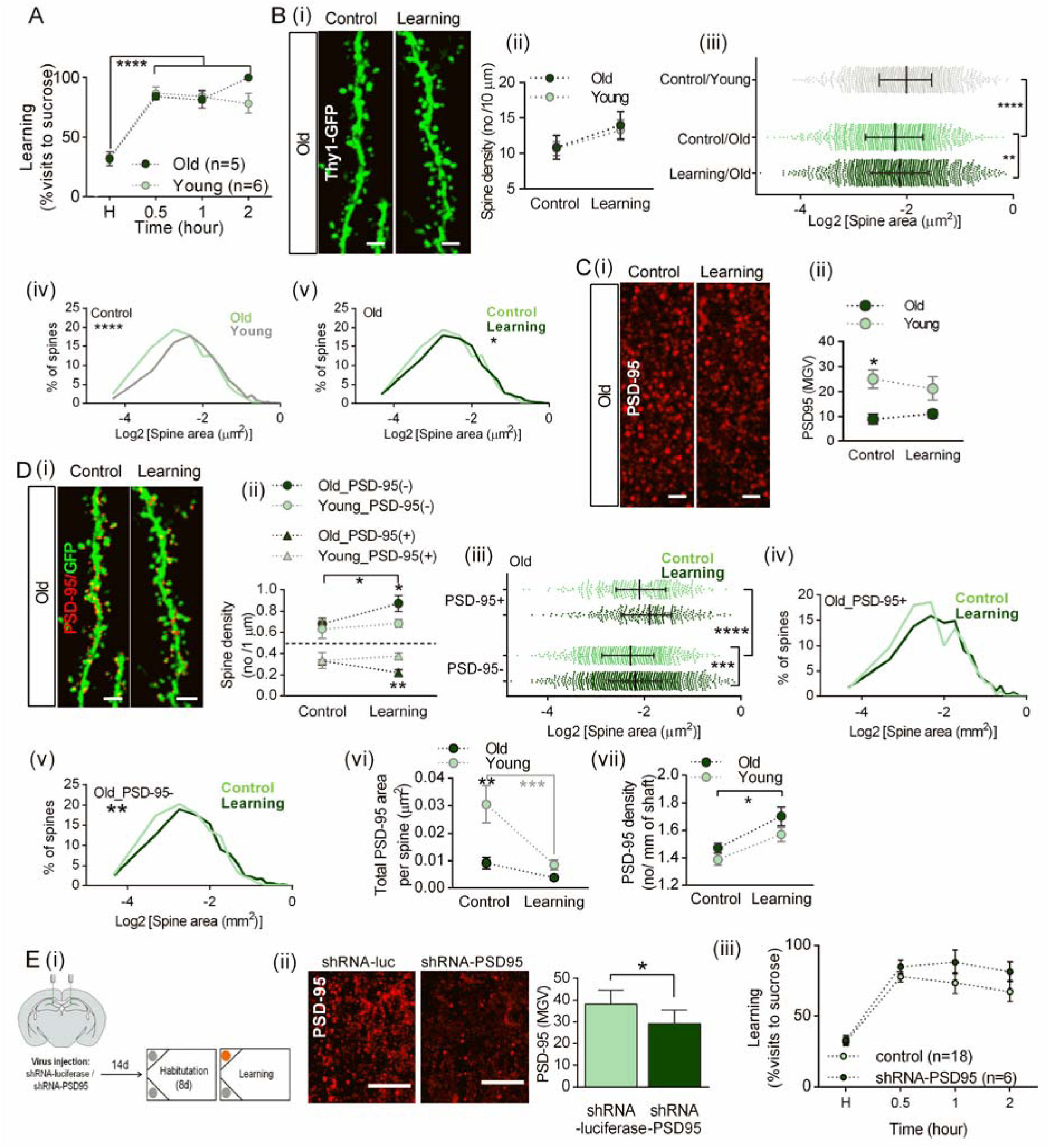
Learning-induced remodeling of dendritic spines in old mice. **(A)** Mean+/-SEM preference for the reward corner during training of old (20±1 month old) and young, adult (5±1 month old) Thy1-GFP(M) mice. Both for young and old mice, the preference of the reward corner was higher during training as compared to the habituation (H). No difference in reward corner preference was observed between the old and young Thy1-GFP(M) mice (****p<0.0001, RM ANOVA with Tukey’s multiple comparisons test; time: F_3, 32_ = 51.88, p < 0.0001; age: F_1, 32_ = 1.037, p = 0.3162). All data for the young mice are the same as in Fig. 1. **(B) (i)** Representative high resolution images of dendritic fragments of the old, control and trained Thy1-GFP(M) mice, scale bars: 2 µm. **(ii)** Mean±SEM linear density of dendritic spines in old and young mice. Linear density of dendritic spines was not affected by the training. There was no difference in spine density in old and young animals (old control mice, n = 4, old learning, n = 5, two-way ANOVA, age: F_1, 16_ = 0.077, p = 0.785; training: F_1, 16_ = 3.401, p = 0.084). **(iii)** Dendritic spines of old control mice were smaller than spines of young control mice, and old trained animals (*****p<0.0001, **p<0.01, Kruskal-Wallis test with Dunn’s multiple comparisons test, H = 40.76, p<0.0001). Graph shows data for individual spines (control, n=972; learning, n=1327) with medians and interquartile range. Scales are Log_2_-tranformed. All statistical tests are performed on non-transformed data. (**iv**) Distribution of spines’ areas of old control mice is shifted toward smaller values as compared to young control mice (****p<0.0001, Kolmogorov-Smirnov test. (**v**) In old mice distribution of the spines’ areas shifted toward larger values after training (*p=0.015, Kolmogorov-Smirnov test). **(C)** (**i**) Representative high resolution images of PSD-95 immunostaining in old, control and trained mice, scale bars: 2 µm. (**ii**) Mean gray value of images was lower in old control mice as compared to young control animals, and it did not differ from the old, trained mice (*p < 0.05, two-way ANOVA with Tukey’s multiple comparisons test, age: F_1, 16_ = 13.59, p = 0.0020; training: F_1, 16_ = 0.0597, p = 0.810, interaction: F_1, 16_ = 0.6769, p = 0.4227). **(D) (i)** Representative high resolution images of PSD-95 immunostaining co-localized with dendritic fragments of the old, control and trained Thy1-GFP(M) mice, scale bars: 2 µm. (**ii**) Average±+/-SEM density of the spines with (+) and without PSD-95 (-). Spines without PSD-95 are more frequent than spines with PSD-95. Training increased density of PSD-95(-) spines in old mice (*p<0.05), and they have more PSD-95(-) spines than young trained mice (*p<0.05, two-way ANOVA with Tukey’s multiple comparisons test, training: F_1, 13_ = 3.532, p = 0.082; age: F_1, 13_ = 3.258, p = 0.0943). After training old mice have less PSD-95(+) spines than young trained mice (*p<0.05, two-way ANOVA with Tukey’s multiple comparisons test, training: F_1, 13_ = 0.7914, p = 0.3898; age: F_1, 13_ = 4.734, p = 0.0486). (**iii**) In the control group, PSD-95(+) spines are bigger than PSD-95(-) spines (****p<0.0001). Only PSD-95(+) spines grew after training (****p<0.0001, Kruskal-Wallis ANOVA followed by Dunn’s multiple comparisons tests, H = 50.62) (control PSD-95-, mice/ spines = 4/ 641; learning PSD-95-, mice/ spines = 5/ 1077; control PSD-95+, mice/ spines = 4/ 329; learning PSD-95+, mice/ spines = 5/ 289). (**iv**) Distribution of PSD-95(+) spines’ areas of old mice did not change after training (p>0.05, Kolmogorov-Smirnov test, D = 0.094). (**v**) The distribution of PSD-95(-) spines’ areas shifted to bigger values after training (**p<0.01, Kolmogorov-Smirnov test, D = 0.087). **(vi)** Average area of PSD-95 immunostaining per spine was smaller in old control mice, as compared to young control animals, and it was not affected by the training (**p<0.01, two-way ANOVA with Tukey’s multiple comparisons test, age: F_1, 13_ = 15.02, p = 0.0019, training: F_1, 13_ = 14.34, p = 0.0023). (**iv**) Average density of PSD-95 puncta in dendritic shaft increased after training both in young and old mice (*p<0.05, two-way ANOVA with Tukey’s multiple comparisons test, age: F_1, 16_ = 3.707, p = 0.0721, training: F_1, 16_ = 13.74, p = 0.0019). **(E) (i)** Experimental timeline. (**ii**) Representative images of PSD-95 immunostaining in CA1 area after overexpression of shRNA for PSD-95 and control shRNA designed for luciferase (*p < 0.05, t-test, t(9) = 2.292), scale bars: 10 μm. **(iii)** CA1 area-targeted shRNA designed for PSD-95 did not affect preference for reward corner in old mice (RM Two-way RM ANOVA, virus: F_1, 23_ = 2.551, p = 0.1239, time: F_3, 69_ = 19.87, p < 0.0001, interaction: F_3, 69_ = 0.3756, p = 0.7709). H, preference of the corner during the last day of the habituation.

Neither age nor training affected density of dendritic spines (**Fig. 3B.i-ii**). However, the medial of dendritic spines’ areas in old control mice was significantly smaller spines than young controls (**Fig. 3B.iii**). Moreover, behavioural training resulted in increased median value of dendritic spines’ areas in old animals (**Fig. 3B.iii)**. These changes were also observed as shifts in distribution of values of dendritic spines’ areas. Distribution of spines’ areas of old control mice was shifted toward smaller values as compared to young control mice (**Fig. 3B.iv**). In old mice distribution of the spines’ areas shifted toward larger values after training (**Fig. 3B.v**).

Intensity of PSD-95 immunostaining in the area CA1 was decreased in the control old mice as compared to the control young group, and it was not changed by the training (**Fig. 3C**).

Next, we calculated density and size of dendritic spines with and without PSD-95 protein. As in young mouse, in old animals PSD-95(-) spines were more frequent than PSD-95(+) spines (**Fig. 3D.ii**). However, in old mice density of PSD-95(-) spines increased after training over the values observed in young trained mice. At the same time point the density of PSD-95(+) spines was lower in old mice as compared to the young animals (**Fig. 3D.ii**). The analysis of the areas of spines with and without PSD-95 protein showed that the spines with PSD-95 have higher median values than spines without PSD-95 (**Fig. 3D.iii**). Moreover, the median values of PSD-95(-) spines’ areas increased after training while the median values of PSD-95(+) spines’ areas did not change (**Fig. 3D.iii**). The change of PSD-95(-) spines in old mice was also observed as a shift of size distribution toward bigger values in learning mice as compared to the controls (**Fig. 3D.v**). No statistically significant change in distribution of PSD-95(+) spines’ areas was observed (**Fig. 3D.iv**). We also analysed PSD-95 puncta to find that the total area of PSD-95 puncta per spine in PSD-95(+) spines was lower in the old control mice, as compared to the young, control animals and it was not affected by the training (**Fig. 3D.vi**). The density of PSD-95 puncta in the shaft increased after training, both in young and old mice (**Fig. 3D.vii**).

In summary, our data indicate age-and spine type-specific remodelling of dendritic spines during memory formation. In young mice training resulted in shrinkage of big spines containing PSD-95. In old mice, the density of PSD-95(-) spines increased, suggesting removal of PSD-95 protein that was presumably translocated to the shaft. At the same time the average size of the spines without PSD-95 increased.

Next, to test the function of PSD-95 protein in old mice, we used lentiviruses encoding shRNA targeted to PSD-95 mRNA (LV:αCaMKII-shRNA_PSD-95-GFP) or luciferase ^8^ (**Fig. 2B**). 20±1 month old, C57BL/6J mice had LVs stereotactically injected into dorsal area CA1, to downregulate PSD-95 expression, and 14 days later they were trained in the IntelliCages (**Fig. 3E.i-ii**). ShRNA for PSD-95 did not impair mice performance neither during initial 30 minutes of the training, nor later (**Fig. 3E.iii**). Since downregulation of PSD-95 in CA1 of old mice increased activity of the mice (extended data, **Fig. 3-1.B**), we also analysed the preference of the reward corner in 10-visit bins (to make the number of learning trials equal between the experimental groups) to find similar effects as described for the time bins (extended data, **Fig. 3-1.B**).

## Discussion

In the current study we analyzed the molecular mechanisms of reward location memory. We show that formation of memory in young adult mice is accompanied by elimination of PSD-95 protein from large dendritic spines and dendritic spine shrinkage in the *stratum radiatum* of CA1 area. Using molecular manipulations *in vivo* we demonstrate that autophosphorylation of CaMKII and CaMKII-dependent destabilization of PSD-95 at the synapse by phosphorylation of PSD-95:serine 73 accelerates memory formation, as the speed of learning is compromised by overexpression of phosphorylation-deficient form of PSD-95:S73A. Long-term downregulation of PSD-95 decreases precision of reward location. In old mice, consolidation of reward location memory results in increased population of spines without PSD-95 and dendritic spines in this category grow. Overall, our data indicate that dynamic regulation of PSD-95 at the synapse is a mechanism for memory formation and stabilization that operates in young animals, but is impaired in aged mice. Thus in old age PSD-95-independent processes underlie learning.

We trained mice to find sucrose reward in one of two active corners of the IntelliCages ^34^. The system allowed for on-line monitoring of mice performance during the training. Preference to visit reward corner was used as a measure of reward location memory and its precision. Both young and old mice increased preference of the rewarded corner during initial 30 minutes after reward location. In young mice the training resulted in shrinkage of dendritic spines that contained PSD-95 in *stratum radiatum* of the area CA1. The total number of the spines with PSD-95 was not altered, however, the size of PSD-95 clusters in spines was decreased. In contrast to young animals, reward location training in aged mice affected mostly spines without PSD-95. Their frequency and size were increased. The spines in old control mice where smaller than the spines in control young adults, however, after training they reached similar size. To our knowledge this is the first study that shows spine type-and age-specific downregulation of PSD-95 protein during memory processes. Since PSD-95 protein controls localization of AMPAR at the synapse ^19,20^, AMPAR currents ^8,19,41^, and synaptic plasticity ^9,10^, the morphological and molecular changes we observe in young mice suggest that formation of memory about reward location is accompanied by weakening of CA1 circuit, and this process is impaired in aged mice. This is in agreement with earlier findings showing that formation of memory about spatial location of a novel object temporarily weakens synaptic transmission ^42,43^. To fully validate whether synaptic transmission in CA1 is indeed altered in our model further experiments are needed. Currently, we can, however, conclude that PSD-95 scaffolding is disassembled during memory formation in young mice. To test the role of this process we performed virally-mediated local manipulations of PSD-95 expression.

CA1 area-targeted long-term downregulation of PSD-95 protein in young mice by overexpression of specific shRNA did not affect initial phase of learning, however, later it impaired precision of spatial memory. Young mice with depleted PSD-95 levels showed lower and less stable preference for the rewarded corner as compared with the control group. This finding is in agreement with the earlier studies showing that PSD-95 expression is dispensable for the formation and expression of recent contextual fear memories, but it is essential for their precision ^14,44^. Surprisingly, this function of PSD-95 is impaired in aged mice which show similar precision of reward location memory to young mice, despite lower levels of PSD-95 protein in dendritic spines. The precision of reward location was also not affected in aged animals by further depletion of PSD-95 in CA1 by PSD-95-targeted shRNA. Thus our data indicate that old mice use PSD-95-independent, or possibly CA1-independent, strategy to precisely remember reward location. This hypotheses need, however, further validations.

Previously, it was shown that synaptic stimulation results in CaMKII-dependent mobilization of PSD-95 from dendritic spine ^28^. This process relies on CaMKII-driven phosphorylation of PSD-95 on serine 73 and requires autophosphorylation of CaMKII:T286 ^26,28^. The function of PSD-95:S73 phosphorylation in memory processes was never tested. Here we tested both the role of autophosphorylation of CaMKII:T286 and CaMKII-dependent phosphorylation of PSD-95:S73 in reward location memory. The training of autophosphorylation-deficient CaMKII mutant mice (T286A) and mice with local overexpression of phosphorylation-deficient PSD-95:S73A in the area CA1 indicate that these processes regulate and speed up early phase of learning. Moreover, autophosphorylation of CaMKII, but not phosphorylation of PSD-95:S73, is important for precision of reward location memory. Our data are in agreement with many earlier studies showing that autophosphorylation of CaMKII, as a key regulator of synaptic plasticity ^29,45,46^ and morphology of dendritic spines and PSDs ^47,48^, also controls formation and flexibility of spatial and contextual memory ^29,48–50^. We demonstrate, however, for the first time the role of CaMKII-dependent phosphorylation of PSD-95:S73 in memory.

Overall, our data show that in young animals learning about spatial location of reward induces elimination of PSD-95 protein from dendritic spines of CA1. Fast learning requires autophosphorylation of CaMKII and CaMKII-dependent phosphorylation of PSD-95 at serine 73. The precision of memory, but not the speed of learning, is sensitive to long-term downregulation of PSD-95 protein levels. Surprisingly in aged animals, this function of PSD-95 is not preserved, as depletion of PSD-95 does not affect precision of memory in old mice. We therefore conclude that in the aged animals, that have no signs of cognitive decline, alternative mechanisms support successful and precise memory formation.

## Acknowledgments, Funding and Disclosure

This work was supported by a National Science Centre (Poland) (Grant No. 2013/08/W/NZ4/00861 and 2015/19/B/NZ4/02996) to KR. AC, MŚ and KR designed the experiments; AC, MŚ, AN, MB, KŁ, RP, MŻ and KR performed and analysed the experiments; AC and KR wrote the manuscript. Authors report no financial interests or conflicts of interest.

## Extended Data

**Figure 1-1.**
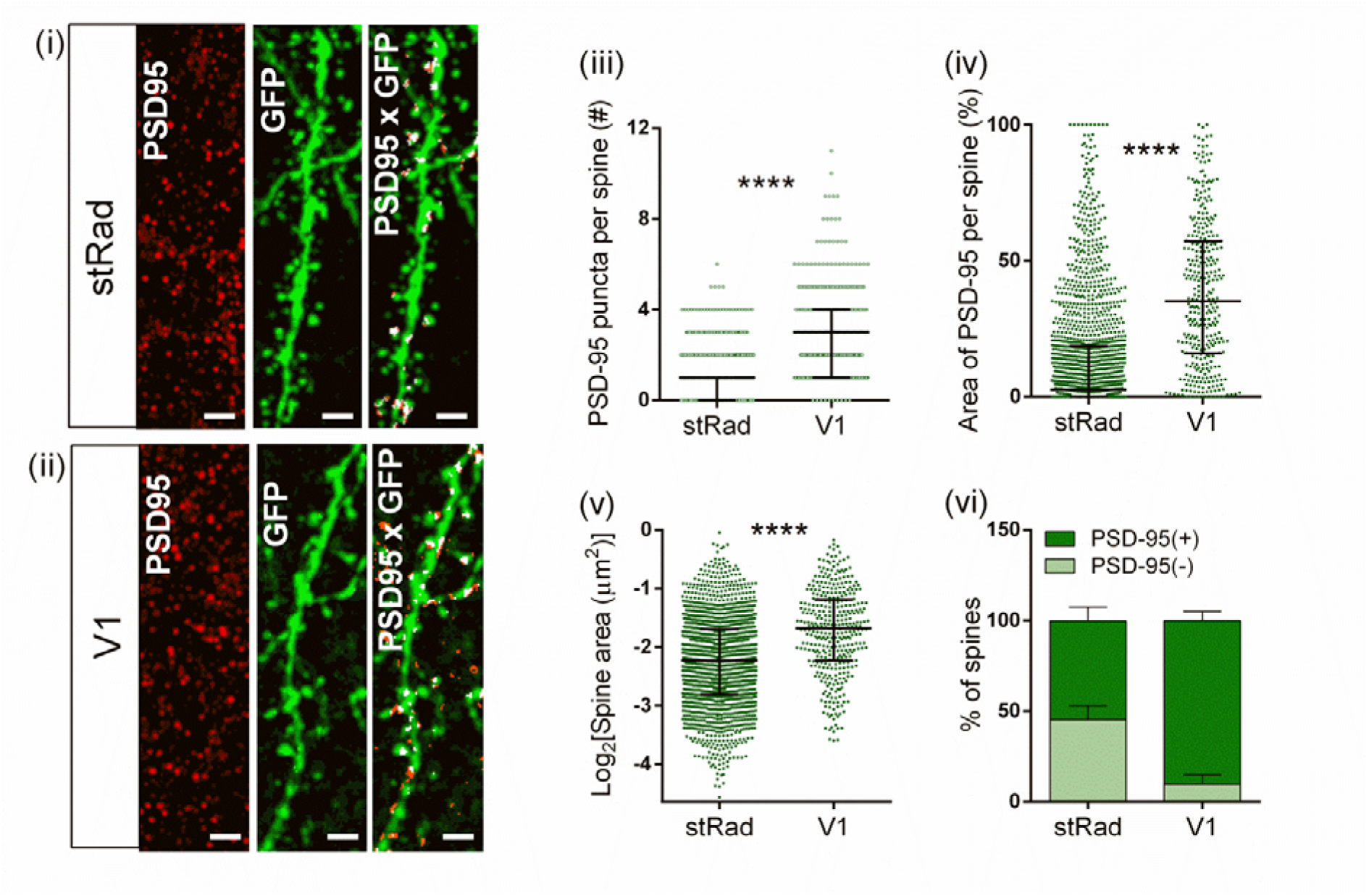
The comparison of PSD-95 protein expression and dendritic spines in the area CA1 and visual cortex (V1). Representative high resolution images of PSD-95 immunostaining, dendritic fragments and co-localization of both in **(i)** *stratum radiatum* CA1 (stRad) and **(ii)** visual cortex (V1) of the control Thy1-GFP(M) mice, scale bars: 2 µm. (**iii**) There are more PSD-95 puncta per dendritic spine in V1, as compared to stRad (****p<0.0001, Mann-Whitney test). **(iv**) Total fraction of PSD-95 immunosignal per spine is higher in V1 than stRad (***p<0.001, Mann-Whitney test). (**v**) Dendritic spines in V1 are bigger than spines in stRad (****p<0.0001, Mann-Whitney test). **(iii-v)** Each dot represents individual spine. (**vi**) There is higher frequency of spines with PSD-95 in V1 than in stRad.

**Figure 2-1.**
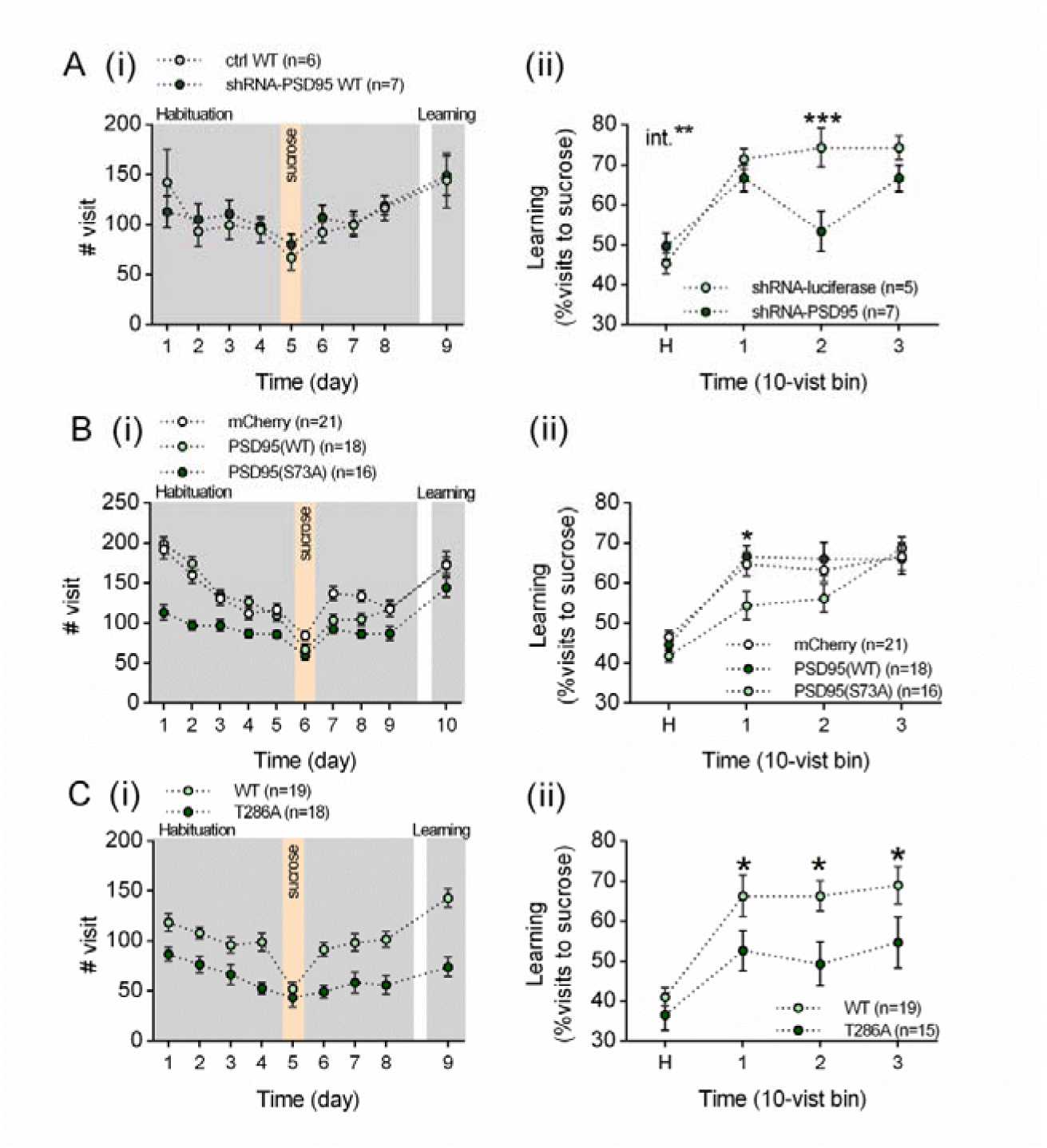
Mice activity during spatial training in the IntelliCages – the effect of modification of PSD-95 and CaMKII. **(A)** Mice behavior after downregulation of PSD-95 protein in the area CA1 by shRNA for PSD-95 in young mice. (**i**) shRNA for PSD-95 did not affect the general mice activity of the mice, measured as total number of visits in the corners (two-way ANOVA, shRNA: (i) virus: F_1, 99_ = 0.2544, p = 0.615; time: F_8, 99_ = 3.579, p = 0.001). **(ii)** shRNA did not affect preference for reward corner during learning measured as % visit of sucrose in 10-visit bins (***p<0.001, two-way ANOVA with Sidak’s multiple comparisons test, virus: F_1, 12_ = 3.536, p = 0.08; time: F_4, 48_ = 10.93, p < 0.0001; interaction: F_4, 48_ = 3.950, p = 0.007). H – the last day of habituation for all experiments. **(B)** Mice behavior after overexpression of PSD-95(WT) and (S73A) in CA1. **(i)** General mice activity in the experiment measured as total number of visits in the corners. Mice with PSD-95(S73A) were less active than other groups during habituation but not during the training (RM ANOVA with Tukey’s multiple comparisons test, virus: F_2, 691_ = 72.74, p < 0.0001; time: F_9, 691_ = 36.81, p < 0.0001; interaction: F_18, 691_ = 3.127, p < 0.0001). (**ii**) Preference for the reward corner during learning measured as % visits of sucrose in 10-visit bins (*p<0.05, two-way ANOVA with Sidak’s multiple comparisons test, time: F_4, 260_ = 30.80, p < 0.0001; virus: F_2, 260_ = 2.567, p = 0.078). **(C)** Young αCaMKII-T286A mutant mice. **(i)** General mice activity in the experiment measured as total number of visits in the corners. Mutants were less active than wild-type mice (RM ANOVA, genotype: F_1, 283_ = 79.23, p < 0.0001, time: F_8, 283_ = 6.591, p < 0.0001). **(ii)** Preference for reward corner during learning measured as % visits to sucrose corner in 10-visit bins. Mutants had lower preference for the rewarded corner than wild-type mice (*p<0.05, RM ANOVA with Tukey’s multiple comparisons test, genotype: F_1, 169_ = 12.91, p = 0.0004; time: F_4, 169_ = 9.037, p < 0.0001).

**Figure 3-1.**
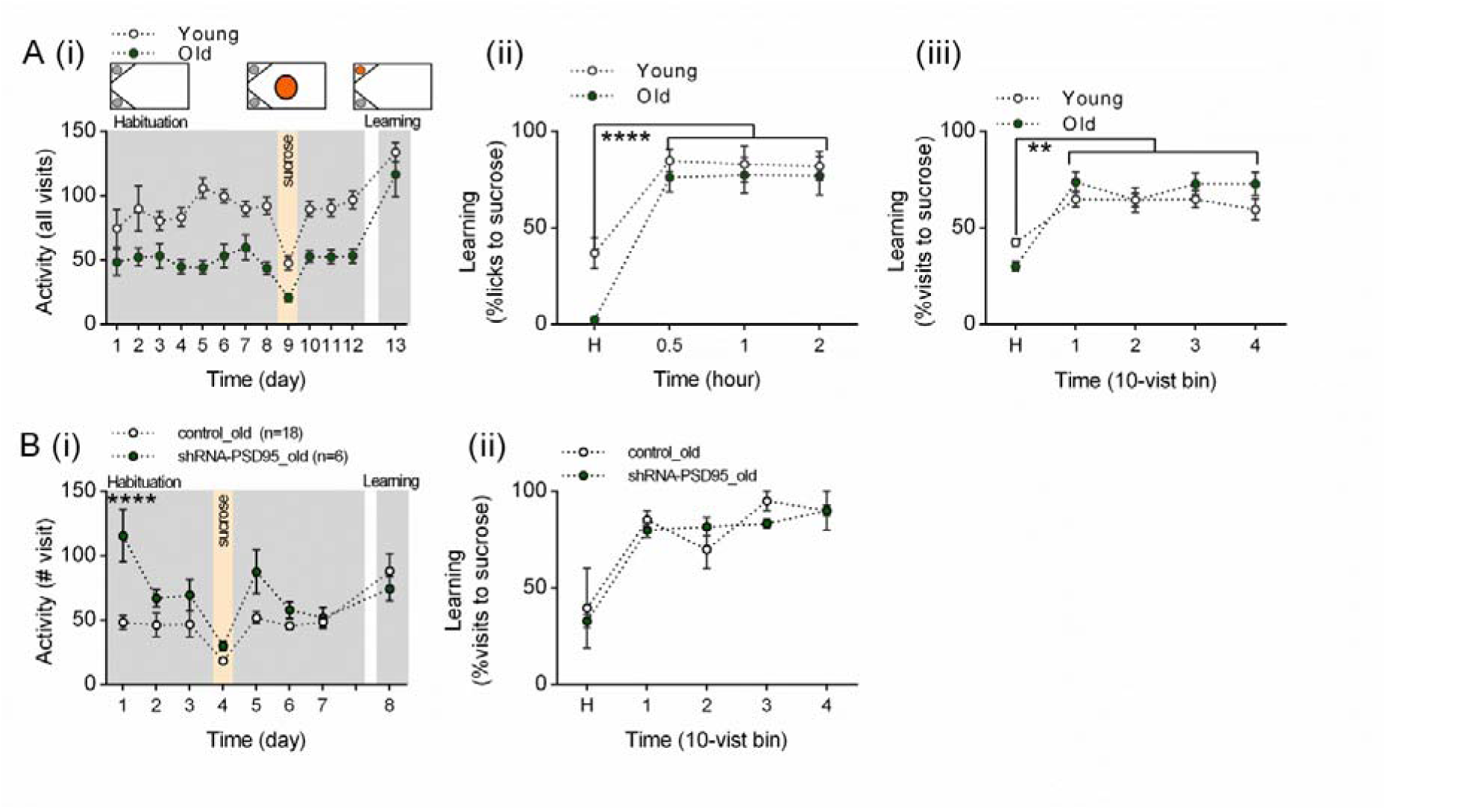
Old mice activity during reward-driven spatial learning in the IntelliCages – the effect of downregulation of PSD-95 in CA1. **(A) (i)** Top, cage setups during training. Gray circles, bottles with water; orange circles, bottles with sucrose. Bottom, total number of visits. Old mice performed less visits than young animals during the habituation phase, but not during spatial learning (RM ANOVA, age: F_2, 593_ = 18.85, p < 0.0001; time: F_12, 593_ = 25.65 p < 0.0001; interaction: F_24, 593_ = 0.713, p = 0.840). **(ii)** Mean±SEM preference for the reward corner during learning. Both young and old mice prefer to drink sucrose than water. H, the last day of the habituation for all graphs (****p<0.0001, RM ANOVA with Sidak’s multiple comparisons test, age: F_1, 211_ = 18.06, p < 0.0001; time: F_4, 211_ = 38.29, p < 0,0001; interactions: F_4, 211_ = 1.041, p = 0.387). (**ii**) Preference for the reward corner during learning measured as % visits to sucrose corner in 10-visit bins (**p<0.01, two-way ANOVA with Sidak’s multiple comparisons test, time: F_4, 208_ = 18.36, p < 0.0001; age: F_1, 208_ = 1.459, p = 0.2285). **(B)** Old mice behavior after downregulation of PSD-95 protein in the area CA1 by shRNA for PSD-95 in young mice. **(i)** shRNA-driven downregulation PSD-95 increased activity of old during habituation but not training (****p<0.0001, RM two-way ANOVA with Sidak’s multiple comparisons test, shRNA: (i) virus: F_1, 19_ = 4.440, p = 0.0486; time: F_7, 133_ = 12.3, p < 0.0001; interaction: F_7, 133_ = 4.987, p < 0.0001). **(ii)** shRNA did not affect preference for reward corner during learning measured as % visit of sucrose in 10-visit bins (***p<0.001, two-way ANOVA with Sidak’s multiple comparisons test, virus: F_1, 6_ = 0.1930, p = 0.6758; time: F_4, 24_ = 39.68, p < 0.0001; interaction: F_4, 24_ = 1.579, p = 0.2120). H – the last day of habituation for all experiments.

